# Nanodelivery of lipids to coral larvae maximises post-settlement survival: Implications for larval ecology and reef restoration

**DOI:** 10.1101/2023.12.20.572712

**Authors:** Nadine M. Boulotte, David Rudd, Peter L. Harrison, Craig Humphrey, Kirsten Benkendorff

## Abstract

Interest in reef restoration is increasing as coral mortality has accelerated at an unprecedented rate. However, high mortality rates of coral early-life stages represent a population bottleneck, which directly impacts the effectiveness of restoration projects. While most coral larvae are considered lecithotrophic and catabolise maternally transmitted lipids to meet metabolic demands; here we demonstrate that coral larvae can be facultative feeders. We used nanoparticles to deliver triacylglycerides to aposymbiotic larvae which resulted in a 30% increase in larval energetic lipids, and a 46% increase in survival rate, up to 16 weeks post-settlement. Changes in phospholipid molecular species in the larvae suggest phagocytosis of the nanoparticles, and an increase in free fatty acids indicates lipolysis of the phagocytosed triacylglycerides. We suggest that a continuum of nutritional strategies should be recognised in coral early-life stages, and that nanoparticles can be used by restoration practitioners to deliver nutritional resources to maximise restoration outcomes.

## INTRODUCTION

Coral reef ecosystems continue to face an unparalleled worldwide decline in coral biomass and diversity due to anthropogenic environmental changes ^1,2^. From 2014 to 2017, researchers around the world observed the most severe, widespread, and longest-lasting global-scale coral bleaching event recorded to date; causing one or more mass bleaching episodes on up to 70% of coral reefs worldwide ^3^. This bleaching event resulted in unprecedented coral mortality, a transformation of the composition of coral assemblages ^4^, a significant decline in larval recruitment on the Great Barrier Reef ^5^; and also contributed to changes in reproductive synchrony in some species ^6^. These accelerating global impacts have catalysed increasing interest and investments in restoration-focused interventions using sexually-produced corals, in the hope of restoring sufficient corals to maintain ecosystem function and services ^7,8^.

Sexual coral propagation consists of collecting coral spawn from multiple breeding colonies for cross-fertilisation and larval culture to produce a genetically diverse population of larvae. Once competency is reached, larvae can be settled onto substrata, and grown in a protected nursery prior to deployment onto restoration sites ^8^; alternatively, larvae can be directly settled onto reef restoration sites ^9^. New technologies are now being considered to assist coral population recovery and adaptation to current and future environmental changes, such as selective breeding and microbiome manipulation (also referred to as “assisted evolution”), epigenetic modifications through preconditioning, promoting long-range genetic exchange (“assisted gene flow”), and genetic engineering ^10–15^. While these approaches are targeting adult corals and coral breeding, fewer interventions are targeting early-life stages; hence why coral early-life stages are now identified as a high-priority knowledge gap in coral restoration interventions ^16^.

Coral early-life stages generally exhibit up to 99% post-settlement mortality during their first year of life in the natural environment ^9,17–19^. Therefore, the post-settlement stage represents a critical population bottleneck, which can limit the effectiveness of sexual coral restoration projects ^16^. While several causes of post-settlement mortality have been investigated over the past decade, lipid depletion resulting from the metamorphosis of larvae into settled polyps, and initiation of calcification ^20^, has been overlooked. Stored lipids play essential roles in marine planktonic larvae as endogenous sources of energy during dispersal and settlement, and can represent up to 70% by weight in coral larvae ^21^. Most broadcast spawning coral larvae are aposymbiotic (i.e., non-symbiotic), and it is generally assumed that coral larvae are lecithotrophic (i.e., non-feeding); therefore, they acquire energy endogenously from maternally allocated lipid reserves for larval development, survival, metamorphosis and settlement ^22–25^. Coral larvae can obtain a considerable amount of energy from endogenous wax esters and triacylglycerides (Boulotte *et al.*, under review), with the former being slowly consumed during embryogenesis, while the latter can be rapidly hydrolysed for short-term energy needs ^21,26^.

The concept of ‘facultative feeding’ (reviewed in ^27^), recognises a continuum of nutritional strategies, rather than the traditional dichotomy between planktotrophy and lecithotrophy in marine invertebrate larvae. Under Allen and Pernet’s ^27^ definition, facultative feeding larvae represent “those larvae that can ingest and digest particulate food, and benefit from the digestion of this food, but do not require exogenous food to complete development to metamorphosis”. Reports of larval feeding remain rare among corals, and it is unclear whether coral larvae could potentially acquire exogenous lipid to supplement their reserves and thus increase their fitness during metamorphosis and settlement. In this study, we hypothesised that coral larvae can be facultative feeders, and that, an exogenous supply of energetic lipids (i.e. triacylglycerides) can boost maternal energy provisions and increase post-settlement survival.

With the aim to reduce the post-settlement population bottleneck and increase coral restoration outcomes, we present a novel approach using emulsion-based nanoparticles to deliver marine triacylglycerides at the larval stage. Untargeted lipidomics using ultra-high-performance liquid chromatography/quadrupole-time-of-flight tandem mass spectrometry (UPLC/QToF-MS/MS) was undertaken to provide an in-depth characterisation of lipid uptake and metabolism in coral larvae exposed to the nanoparticles.

## MATERIALS AND METHODS

### Coral spawning and larval rearing

Four gravid colonies of *Acropora tenuis* were collected from Falcon Island (Central Great Barrier Reef - 18.7694°S, 146.5329°E, permit number: G12/35236.1) in October 2018 and transferred to temperature-controlled, flow-through aquaria at the Australian Institute of Marine Science’s National Sea Simulator facility for spawning. Spawning occurred four nights after the full moon, on the 24/10/2018 around 6:30 pm. Larval rearing took place in 500 L larval rearing tanks supplied with temperature-controlled, flow-through filtered seawater (25 L/h) inside a temperature-controlled room set at 27 °C. When embryogenesis was complete, and early stages of larvae had developed (∼24 h post-fertilisation), aeration was supplied, and seawater flow rate was increased to 50 L/h. Larval rearing was continued until larvae reached settlement competency.

### Nanoparticle preparation

Nanoparticles (NP) were prepared using a Lecithin-in-Water emulsion method as described in Yanasarn and collaborators ^28^. Briefly, 400 mg of L-α-Lecithin (Sigma^©^) was dissolved in 100 mL of warm MilliQ water. The mixture was heated to 55 ℃ with continuous stirring until a milky consistency was formed. While maintaining agitation, either 400 mg or 800 mg of marine triacylglycerides (Blackmores^®^ fish oil) was added to the dissolved lecithin; after which, a 2% v/v solution of Tween-20 was added drop-by-drop. The heater plate was turned off to allow the mixture to slowly cool with continuous stirring. The NP mixture was added to the experimental containers once it had reached room temperature. Three different NP treatments were prepared: 1) Lecithin emulsion without the addition of marine lipids, referred as NP(0), 2) Lecithin emulsion containing 400 mg of marine lipids, referred as NP(400) and, 3) Lecithin emulsion containing 800 mg of lipids, referred as NP(800).

### Experiment design and sample collection

Sixteen 2 L glass jars with aeration were set up; of which 4 replicates were randomly assigned to the Control (larvae only) and each nanoparticle treatment NP(0), NP(400) and NP(800). Once the larvae were competent to settle (4.5 days old), ∼2000 larvae were transferred into the 2 L aerated glass jars containing 1800 mL of FSW; after which, 50 mL of nanoparticles mixture was slowly added to the aerated glass jars. After 8 h of incubation, the seawater from each glass container was replaced, and any fat residue was wiped from the glass jar. One-hour after the water change, ∼500 larvae were taken from each replicate, washed three times under a gentle stream of FSW using a 106 µm mesh filter and transferred into a 2 mL cryovial. Samples were flash-frozen in liquid nitrogen and stored at -80 ℃ until processing (N=16 samples for a total of ∼8000 larvae).

### Coral larvae settlement and recruit survival monitoring

Following the sample collection (approximately 10 h post-NP exposure), the remaining larvae were pooled for each treatment group and transferred into settlement tanks. Larval settlement took place inside four separate 2 L plastic containers (one tank per treatment/control set up with a FSW flow-through system) containing 55 aragonite settlement plugs per treatment. Each plug was previously biologically conditioned for 3 months in an outdoor tank set up with non-filtered flow-through seawater to allow the development of biofilm and crustose coralline algae (CCA) providing natural settlement cues for coral larvae ^29–31^. The settlement tanks (with plugs) were placed inside a large flow-through water bath set to 26.5 °C to help regulate the water temperature inside each tank.

Two weeks post-settlement, high-resolution macro-photographs of settled coral spat on plugs were taken with a Nikon D810 to assess the initial settlement and survival rate. Subsequently, all settlement plugs containing settled spats were labelled, randomly mixed, and transferred to a new tray and tank for longer-term survival monitoring. The recruits were reared in a 250 L semi-open aquarium system receiving two turnovers of new seawater daily, with artificial lighting, media filter and protein skimmer to remove excess nutrients. Lighting levels were maintained at PAR readings of 140-160 µmol m^-2^ s^-1^ with a one-hour ramp from 07:30-08:30 to “full” intensity until a one-hour ramp down from 15:00-16:00, and seawater temperature was maintained at 26.5° C. High-resolution photographs were taken 7-, 12- and 16-weeks post-settlement to monitor the survival rate of coral juveniles over time. Coral recruit survival was assessed by counting the number of plugs containing at least one recruit, during each monitoring period. This method was employed in order to match the restoration aim of one surviving recruit per plug and to avoid losing survivorship data as a result of the fusion of some coral recruits on the plugs over time.

### Lipid extraction of coral larvae

Total lipids were extracted using a modified Folch *et al.,* ^32^ procedure scaled down for coral larvae (Boulotte *et al.,* **bio-protocol**). Briefly, 500 larvae were left to defrost on a 1.5 x 4 cm piece of filter paper, and then placed into a 9 mL glass tube containing 1.5 mL of ice-cold methanol. Larvae were gently removed from the filter paper and homogenised using a Chattaway spatula to facilitate cell breakdown. After which, 3 mL of chloroform and 1 mL of 0.9% NaCl were added, followed by several incubations (4°C) and centrifugation steps to reach phase partitioning. The lower phase layer was collected, filtered through glass wool, and then dried under a stream of nitrogen. The dried lipid extract was quantified gravimetrically on an analytical balance (± 0.0001 g) and resuspended at a concentration of 10 mg/mL in hexane with butylated hydroxytoluene (BHT 50 mg/mL) to inhibit lipid peroxidation ^33^. All lipid extracts were stored under nitrogen at -80 °C prior to lipidomic analyses.

### Targeted and non-targeted lipid profiling

For non-targeted lipid profiling, the extracts were aliquoted and diluted to 1 mg/mL in methanol:isopropanol:ammonium acetate (70:20:10). A separate aliquot was prepared and diluted to 1 mg/ml in chloroform:methanol:ammonium acetate (60:30:10) for the wax ester targeted lipidomic analysis ^34^. For each aliquot, qualitative and semi-quantitative analysis of coral larvae lipid molecules was performed using a MS/MS^ALL^ direct infusion acquisition technique via an ultra-high performance liquid chromatography system (UHPLC, Shimadzu) coupled with a quadrupole time-of-flight mass spectrometer (Triple TOF 5600, AB SCIEX), in both positive and negative ion modes. The instrument was operated using Analyst® 1.5 software. Lipid molecules were identified using Lipidview^TM^ software (Version 1.2, AB SCIEX) with a personalised lipid database containing coral wax ester profiles, and lipid identification was based on chromatographic retention, accurate mass and tandem mass spectrometry (MS/MS^ALL^) fragmentation patterns. Ionised lipids detected with a signal-to-noise ratio (s/n) over 5 were included in the analysis ^35^. The lipid nomenclature was based on the LIPID MAPS recommendations, where O denotes (alkyl/acyl) ether lipids.

### Data visualisation and statistical analyses

To test the null hypothesis that the survival of coral recruits did not differ significantly between larvae treated with NPs and control larvae, the survival data were modelled using the non-parametric Kaplan-Meier survival analyses with the “survminer” ^36^ and “survival” ^37^ packages in Rstudio.

The molecular lipid species compositions of each lipid class were identified in Lipidview^TM^ and semi-quantified based on the peak area normalised to the sum of all peaks in the extract. The lipid profile for each class of lipids was collated and characterised across all extracts using a hierarchical cluster analysis and visualised in heatmaps using the “pheatmap” and “ggplot2” ^38^ packages in Rstudio ^39^. All data visualisations were realised using the following packages: “ggplot2”, “tidyr” ^40^, “RColorBrewer” ^41^ and “Patchwork”.

To test the null hypothesis that the lipid composition of coral larvae did not differ between Control and NP treated larvae, the relative composition of lipid molecules in each replicate extract was calculated using the peak area normalised with respect to total lipid (= sum of all lipids) in each sample. The data were then imported into PRIMER v7 ^42^ with the PERMANOVA + add-on ^43^. Principal coordinate analysis (PCO) plots, with vectors overlay based on a Pearson rank correlation greater than 0.8, were generated using a Bray Curtis similarity matrix between samples – on the square root transformed data – to visualise the relationships between larval treatment groups based on the multivariate lipid compositions in two dimensional space. PERMANOVA tests were performed using type I sums of squares, unrestricted permutation of raw data and 9999 permutations with “Treatment” (4 levels: Control, NP(0), NP(400) and NP(800)) as a fixed factor. A post hoc pair-wise comparisons test among all pairs of levels of ‘Treatment’ was used to identify any statistically significant differences, and a similarity percentages (SIMPER) analysis was used to determine which lipid class contributed the most to the dissimilarity among all pairs of levels of “Treatment”.

## RESULTS

### Survival of ex-situ reared coral recruits following larval feeding

Monitoring of survival of newly settled coral spat on the settlement plugs showed a significant nanoparticle (NP) treatment effect on coral recruit survival rate over time (p < 0.0001, Kaplan-Meier survival estimate, Fig. 1a). Overall, by the end of 16 weeks, there was a 46% increase in survival rate when coral recruits were incubated with either 400 mg or 800 mg of marine triacylglyceride nanoparticles (Fig. 1a and 1b). Two weeks post-settlement, the survival rate of the control recruits was ∼10% lower than both of the NP treatment recruits (89% for the Control, compared with 98% for NP(800) and 100% for both NP(0) and NP(400), Fig. 2). There were no substantial differences in the subsequent survival rates for any of the treatments between two weeks and seven weeks post-settlement. However, at 12 weeks post-settlement, there was a 52% mortality rate among recruits in the Control compared to 32%, 27% and 18% mortality rate in NP(0), NP(400) and NP(800) recruits, respectively. By 16 weeks, only 35% and 46% of the Control and NP(0) recruits survived respectively, compared to a 65% and 64% survival rate observed in NP(400) and NP(800) recruits, respectively (Fig. 1b).

**Figure 1.**
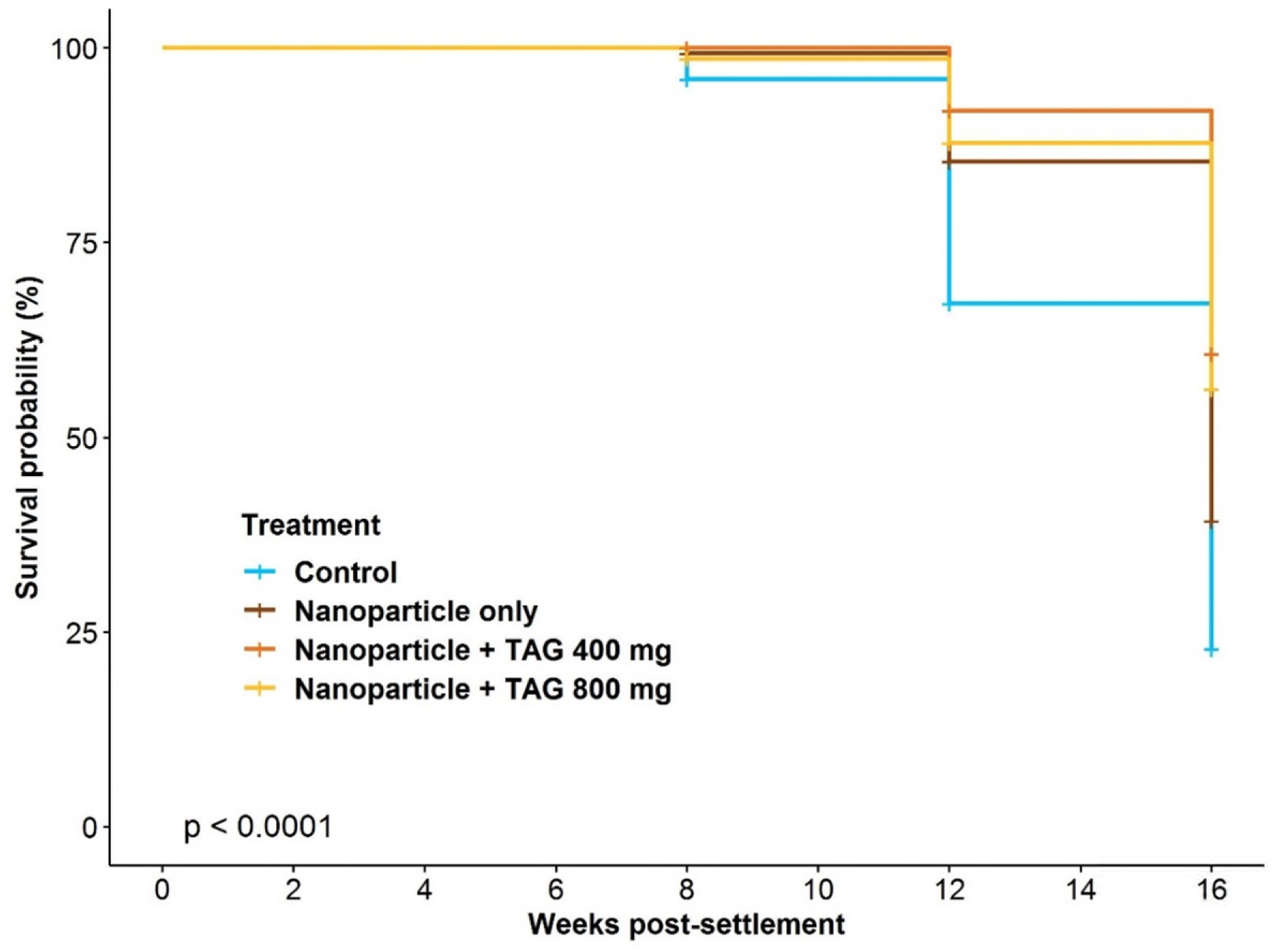
Kaplan-Meier analysis of A. tenuis coral recruit survival on aragonite plugs monitored for 16 weeks post-settlement, following exposure to three levels of nanoparticles for 8 hours at 4.5 days after fertilization, versus control larvae that were not nutritionally provisioned. The lecithin nanoparticles contained 0, 400 or 800 mg of marine triglycerides (TAG).

**Figure 1b.**
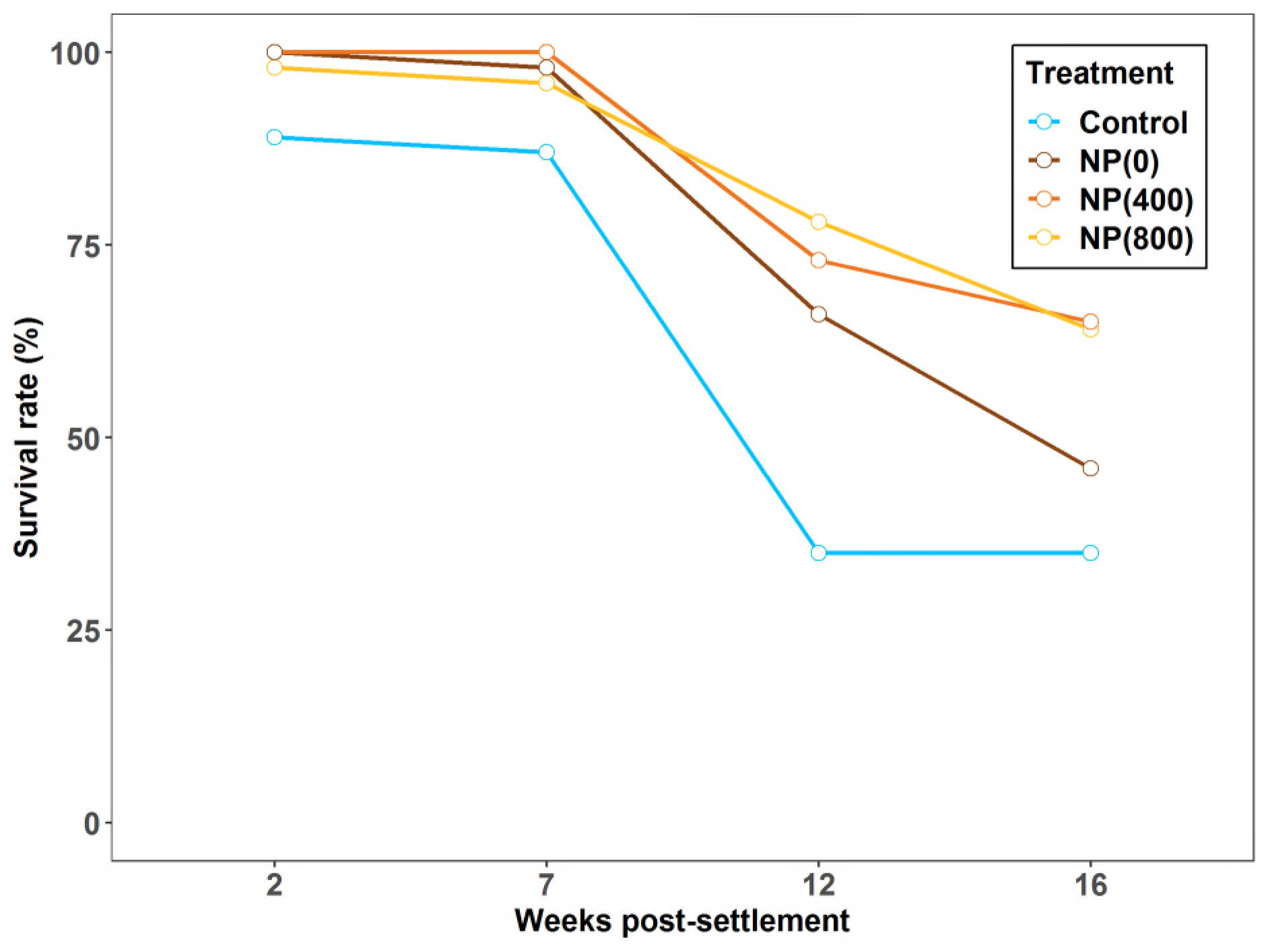
Survival rate of A. tenuis coral recruits on aragonite plugs monitored for 16 weeks post-settlement, following exposure to three levels of nanoparticles for 8 hours at 4.5 days after fertilization, versus control larvae that were not provision. All plugs were randomly placed in the same tank for the 16 weeks survival monitoring (psedoreplication), hence why no error bars are displayed on the graph.

**Figure 2.**
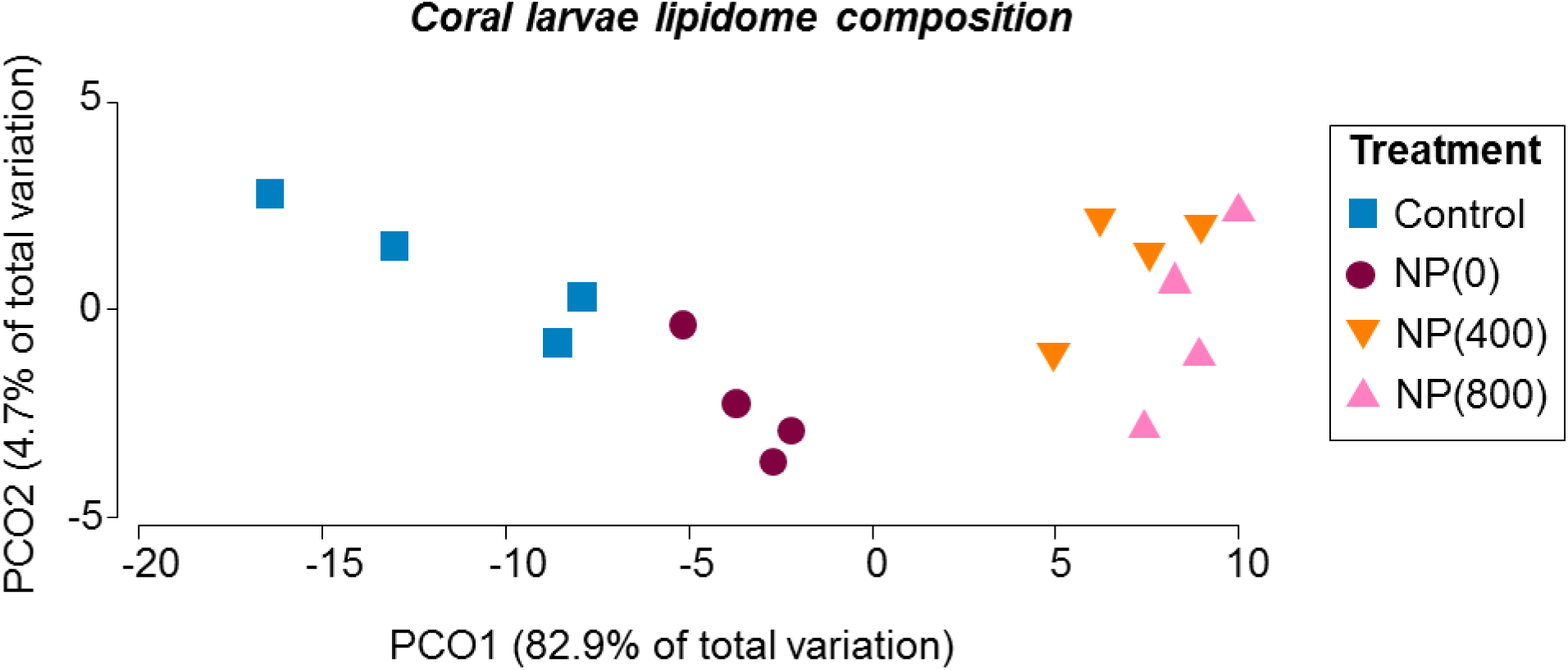
PCO plot of coral larvae lipidome composition generated using a Bray Curtis similarity matrix between samples based on square root transformed data to visualise the separation between larval treatment groups based on the multivariate lipid compositions in two-dimensional space.

### In-depth lipidomic analysis of coral larvae after exposure to nanoparticles

A total of 16 fatty acids were detected in the coral larvae samples, as well as 42 lipid classes including structural lipids (phospholipids), lipids involved in cell signalling (sphingolipids), and energy storage lipids (glycerolipids, sterols, wax esters, free fatty and hydroxyl derivatives of fatty acids, Table S10).

The coral larvae lipidome composition changed significantly after larvae were incubated with nanoparticles (PERMANOVA, p= 0.0001, Table S1, Fig. 2), and the lipid composition of larvae from the nanoparticles treatments were significantly different from the control larvae (Fig. 2, Pair-wise test, Control/NP(400): p=0.0265, Control/NP(800): p=0.0305, and Control/NP(0): p=0.0286, Table S2). This trend was also observed on the principal coordinates analysis plot (PCO) of the entire coral larvae lipidome, where the control larvae were separated from the treatment larvae, showing that some changes occurred in the larvae lipid profiles after incubation with the nanoparticles (Fig. 2).

Significant differences were also observed in the lipidome of larvae exposed to the empty nanoparticles in comparison to those exposed to the marine TAG nanoparticles (Pair-wise test, NP(0)/NP(400): p= 0.0298, and NP(0)/NP(800): p= 0.0316, Table S2). However, there was no significant difference in lipid composition between NP(400) and NP(800) larvae (Pair-wise test, p= 0.4802, Table S2). This trend was also observed in the PCO, where NP(400) and NP(800) larvae are clustered strongly together, highlighting a complete shift in lipidome composition after incubation with nanoparticles containing marine TAG (Fig. 2). For simplicity, detailed lipid results from only Control, NP(400) and NP(800) larvae are displayed in subsequent figures to highlight the differences in lipidome composition arising from exposure to the marine triacylglycerides.

Energetic lipids, structural lipids and fatty acids accounted for 90.5%, 97% and 85.6% respectively of the variation across treatments; with the NP treatments clustered strongly together and separated from the Controls (Fig. 3a, b, c). Significant differences in energy storage lipids, structural lipids and fatty acid profiles were also observed in coral larvae after incubation with nanoparticles (Fig. 3a, b, c, PERMANOVA, p= 0.0029, p= 0.0027 and p= 0.0016 respectively, Table S3, S4 and S5). In contrast, there were no significant changes in lipids involved in cell signalling (Fig. 5d, PERMANOVA, p= 0.1254, Table S5).

**Figure 3.**
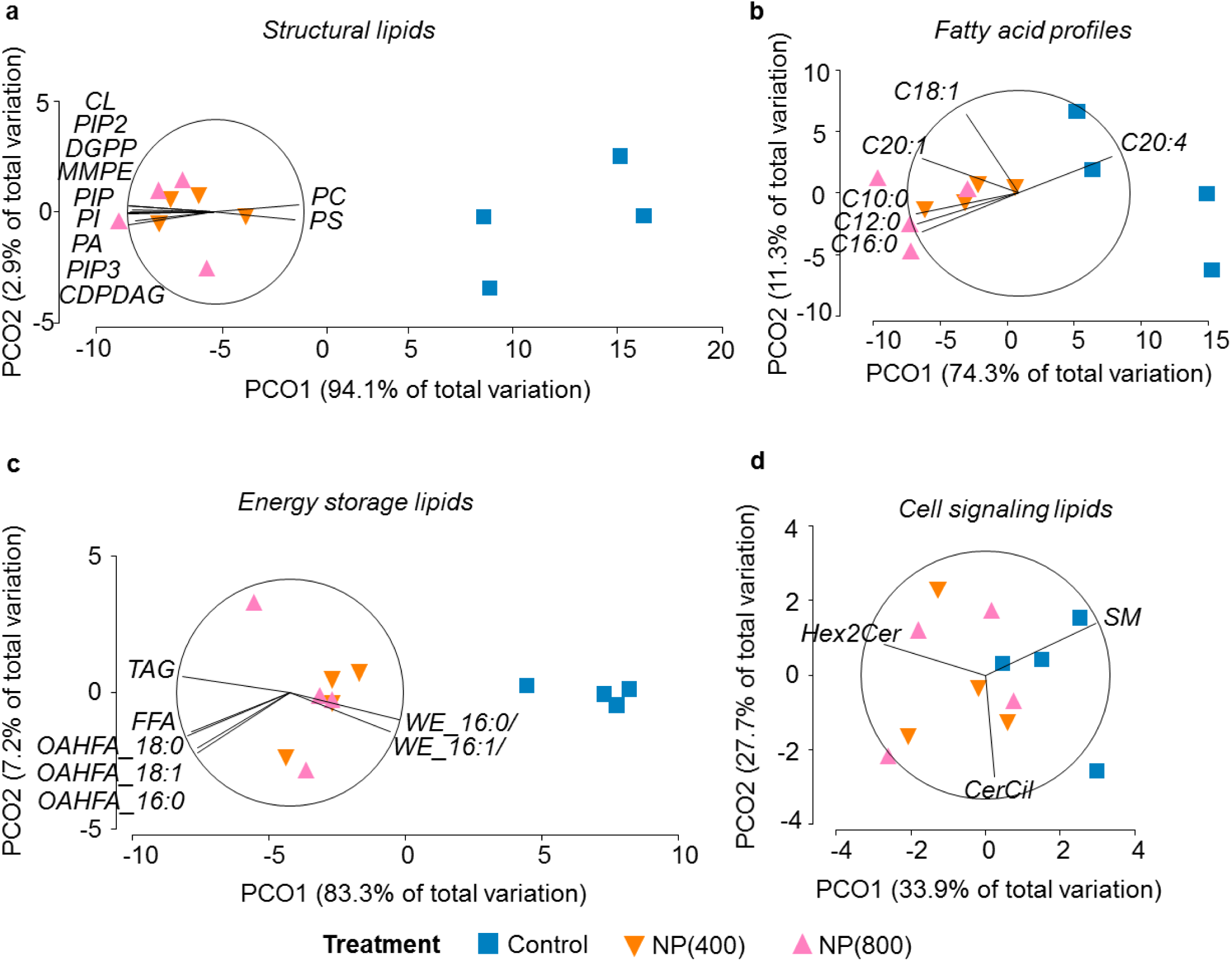
PCO plots of coral larvae lipid categories generated using a Bray Curtis similarity matrix between samples, based on square root transformed data, to visualise the differences in the composition of lipids among treatments. Vector overlays are based on a Pearson rank correlation greater than 0.8. **A**: Structural lipids: Cardiolipin (CL), Phosphatidylinositol 4,5-bisphosphate (PIP2), Diacylglycerol Pyrophosphate (DGPP), Monomethyl-phosphatidylethanolamine (MMPE), Phosphatidylinositol (PIP), Phosphatidylinositol (PI), Phosphatidic Acid (PA), Phosphatidylinositol (3,4,5)-trisphosphate (PIP3), Cytidine Diphosphate Diacylglycerol (CDPDAG), Phosphatidylcholine (PC), Phosphatidylserine (PS). **B**: Fatty acid profiles. **C**: Energy storage lipids: Triacylglyceride (TAG), Free Fatty Acid (FFA), Wax Esters (WE_16:0/, WE 16:1/), the hydroxy fatty acids (OAHFA_18:0, OAHFA_18:1, OAHFA_16:0). **D**: Cell signalling lipids: Sphingomyelin (SM), Dihexosylceramide (Hex2Cer), Ceramide Ciliatine (CerCil).

**Figure 5.**
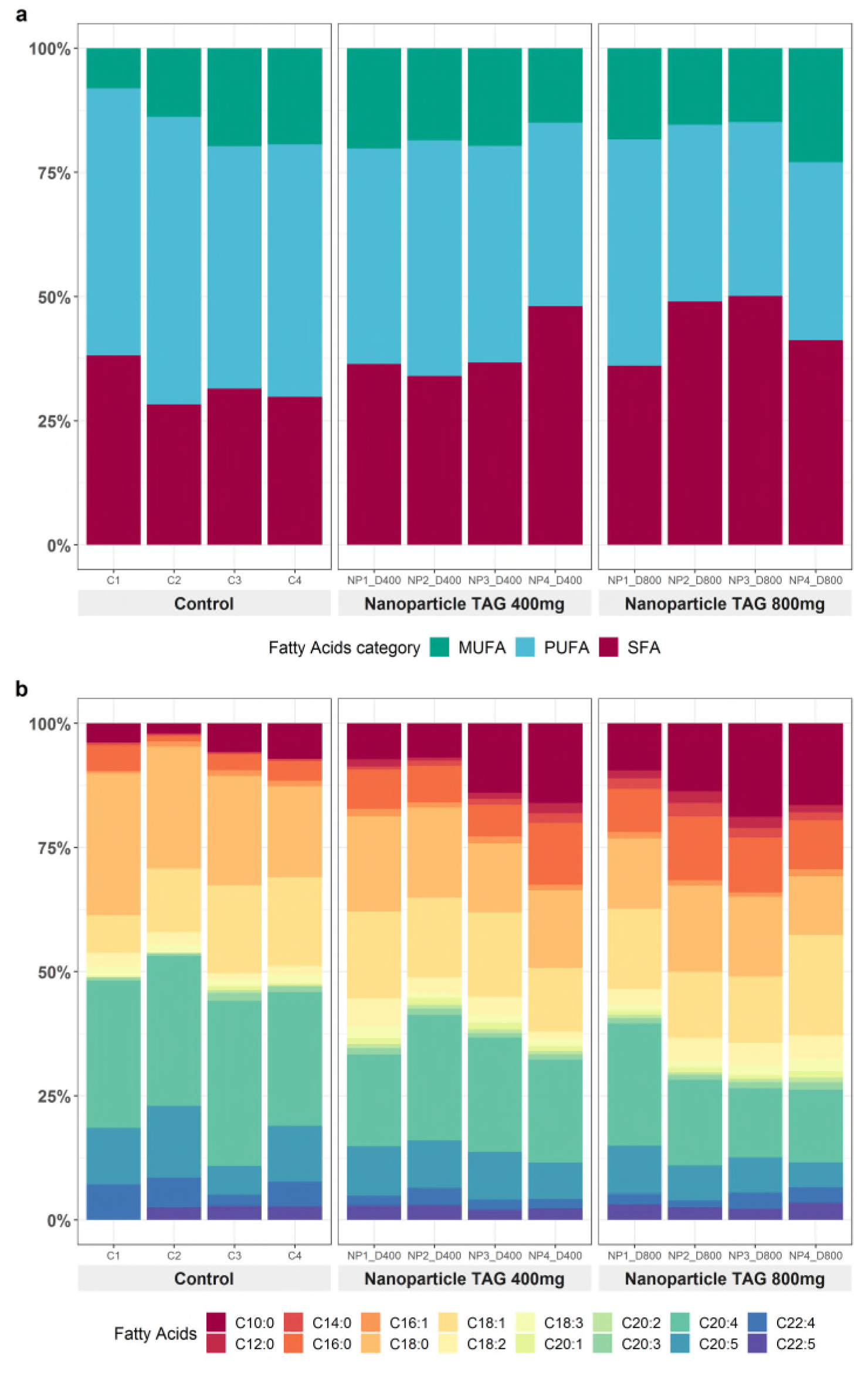
Fatty acid profiles of coral larvae exposed to different nanoparticle treatments. A pool of 500 larvae were analysed for each of the 4 replicates per treatment. Fatty acid composition is presented based on percent peak area relative to total amount of respective lipid class. MUFA = monounsaturated fatty acids, PUFA = Polyunsaturated fatty acid and SFA = Saturated fatty acid.

We observed a 23% and 30% increase in the proportion of energy storage lipids in NP(400) and NP(800) larvae respectively compared to the Control larvae (Fig. 4a). The similarity of percentages (SIMPER) analysis highlighted that the triacylglycerides (TAG), monoalkyl diacylglyceride (MADAG), free fatty acids (FFA), wax esters 16:0 (WE_16:0) and, the hydroxy fatty acids (OAHFAs) were responsible for up to 70% of the dissimilarity in the lipidome composition among treatments and Control (Table S7). For example, we observed an increase in TAG (NP(800): +53%, NP(400): +50%), MADAG (NP(800): +40%, NP(400): +28%), and FFA (NP(800): +90%, NP(400): +88%); while there was a 6% decrease in WE_16:0 in both NP treatments (Fig. 6b). Interestingly, OAHFA_18:0 and OAHFA_18:1 were not detected in the control samples but were present in samples from both NP treatments (Fig. 4b).

**Figure 4.**
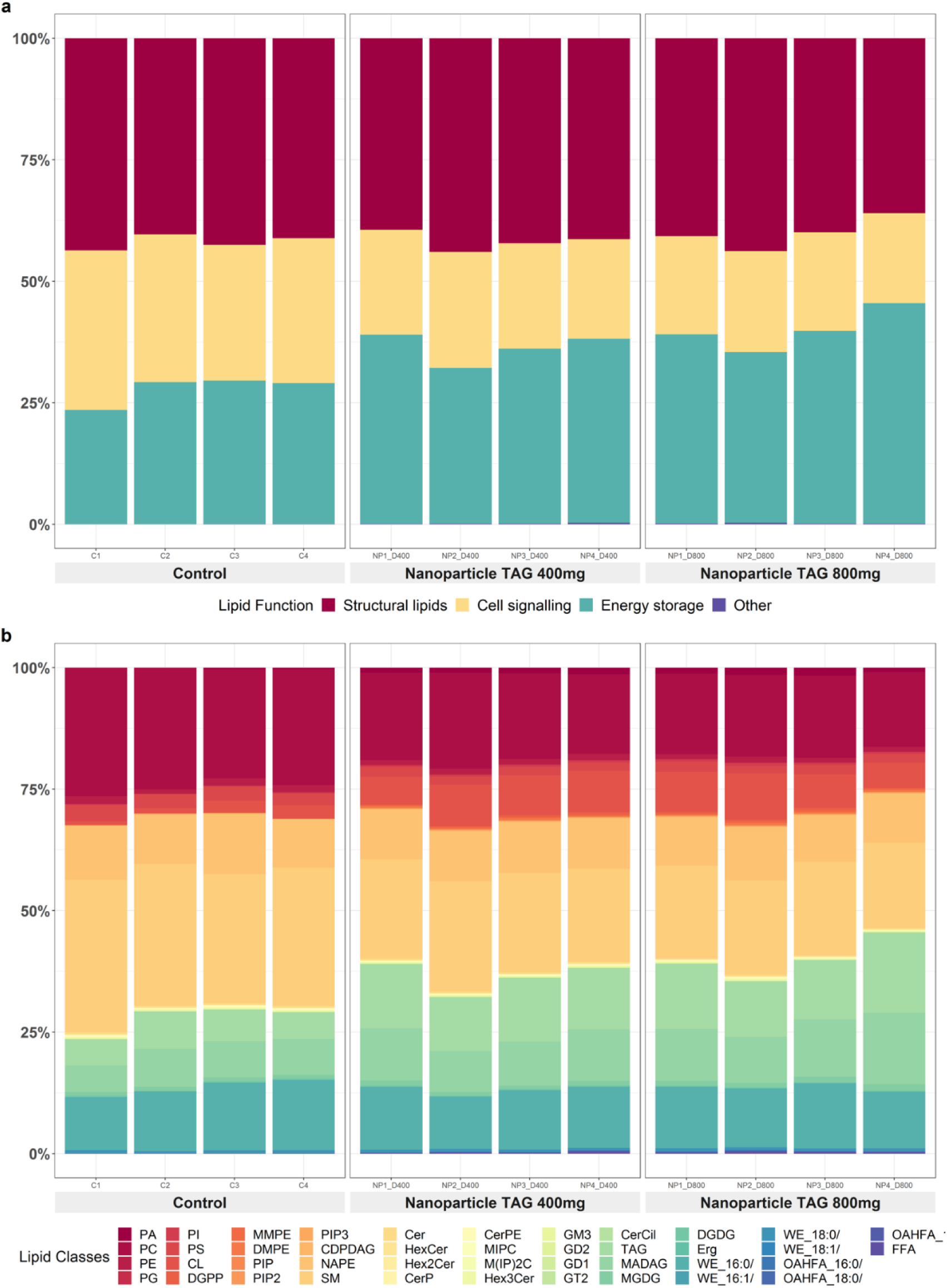
Lipid profiles of coral larvae exposed to the different nanoparticle treatments. A pool of 500 larvae were analysed for each of the 4 replicates per treatment. Lipid data is presented in percent peak area relative to total amount of respective lipid class. The list of the different lipid classes can be found in Table S3.10.

**Figure 6.**
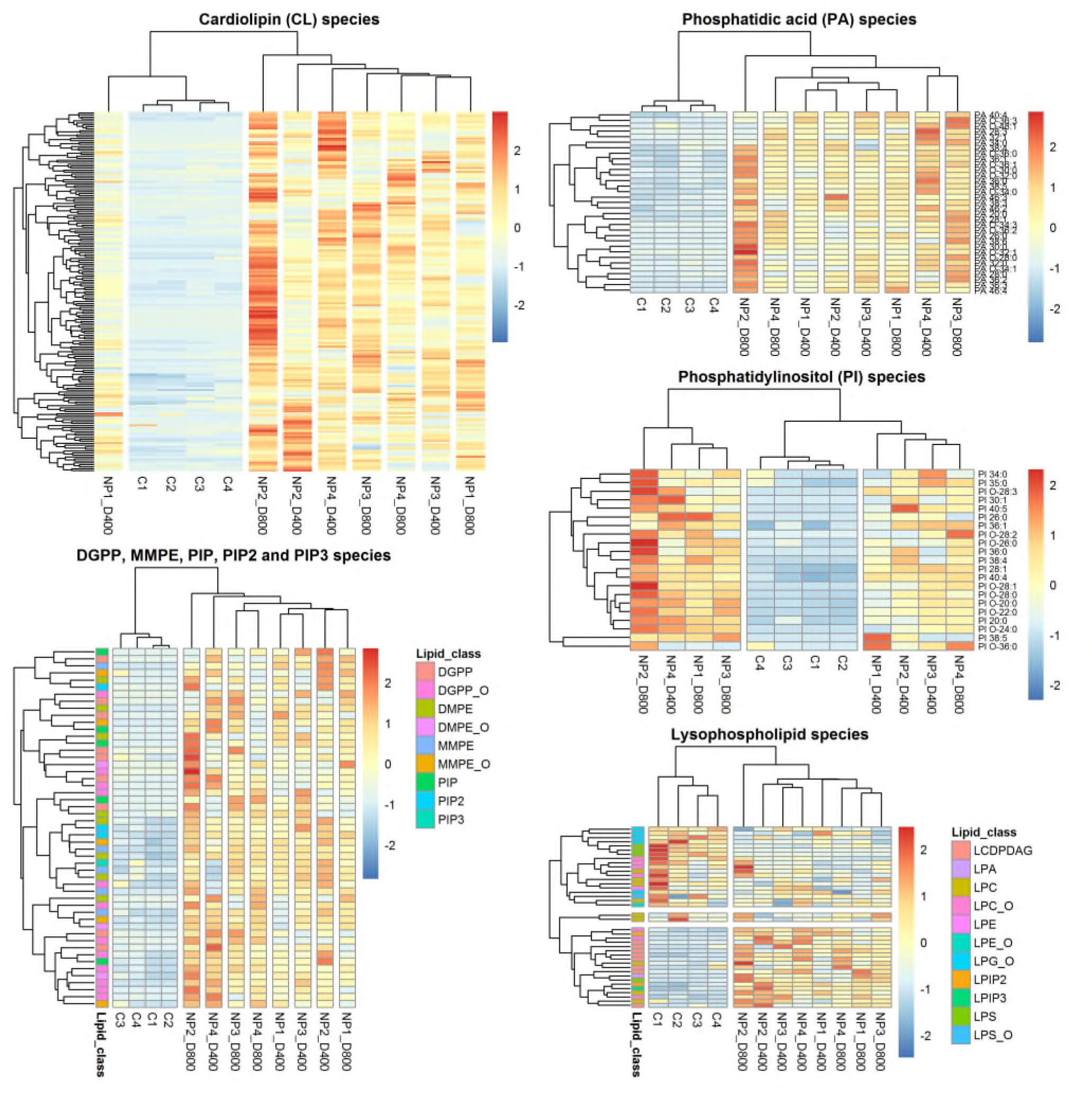
Heatmaps of phospholipids molecular species within each lipid class detected in treatments (NP1 to NP4) and control (C1 to C4). Red and blue correspond to higher and lower relative amounts, respectively. Data were scaled by row.

Within the structural lipids, the SIMPER analysis highlighted that cardiolipin (CL), phosphatidylcholine (PC), phosphatidic acid (PA), phosphatidylinositol (PI), diacylglycerol pyrophosphate (DGPP), and monomethyl phosphatidylethanolamine (MMPE) accounted for up to 70% of the dissimilarity in the lipidome composition among NP treatments and Control (Table S8). There was a greater than 75% increase in CL, >80% increase in PA, >88% increase in PI, and a 93% increase in both DGPP and MMPE across all NP treated larvae; whereas there was a decrease in PC (NP(800): -48%, NP(400): -36%) relative to control larvae (Fig. 4b).

The total fatty acid profiles of NP treated larvae displayed a different composition compared to the control larvae (Fig. 5a, b). We observed an increase in saturated fatty acid (SFA, NP(800): +27%, NP(400): +18%), accompanied with an overall decrease in polyunsaturated fatty acid (PUFA, NP(800): -39%, NP(400): -23%), and a 16% decrease in monosaturated fatty acid (MUFA) in both NP treatments (Fig. 5a). Capric acid (C10:0), lauric acid (C12:0), myristic acid (C14:0), palmitic acid (C16:0), stearic acid (C18:0), arachidonic acid (C20:4), eicosapentaenoic acid (EPA, C20:5), and adrenic acid (C22:4) were responsible for up to 70% of the dissimilarity in the fatty acid composition among NP treatments and Control (Table S9).

Fatty acids that increased in NP treated larvae were capric acid (NP(800): +66%, NP(400): +54%), myristic acid (NP(800): +80%, NP(400): +60%), and palmitic acid (NP(800): +73%, NP(400): +66%). Lauric acid (C12:0) was not detected in control samples but was present in both NP treatment samples (Fig. 5b). In contrast, there was a significant decrease in stearic acid (NP(800): -53%, NP(400): -35%), arachidonic acid (NP(800): -67%, NP(400): - 36%), eicosapentaenoic acid (NP(800): -57%, NP(400): -22%), and 1.5-fold decrease in adrenic acid in both NP treatments compared with the control (Fig. 5, PERMANOVA p= 0.0016, Table S5).

### Dynamics of molecular lipid species in coral larvae after exposure to nanoparticles

The MS/MS^ALL^ analysis identified 1071 molecular lipid species encompassing 9 lipid groups (i.e. phospholipids N=509, ether phospholipids N=126, lysophospholipids species N=27, sphingolipids N=131, glycerolipids N=204, sterols N=2, wax esters N=51, hydroxy fatty acids N=16, and free fatty acid N=5).

Within the structural lipids (i.e. phospholipids, ether phospholipids and lysophospholipids), >70% of molecular species displayed a difference in their relative contribution by either being present in very low abundance or being absent in the Control compared to the NP treatments (Fig. S1). For instance, an increase in abundance, as well as a change in molecular speciation, was observed in CL, PA, PI, PIP, PIP2, PIP3, DGPP, DMPE, MMPE, CDPDAG, PC and their Lyso species; with an explicit Control vs NP treatment clustering highlighted by the hierarchical cluster analysis (Fig. 6). Lipid species belonging to PI were the only type displaying an increased abundance with increasing NP treatment concentration (Fig. 6). CL was the most diverse group with 183 molecular species detected across all samples, of which 88 species were only present in the NP treatments. Of the CL species present in Control, CL 72:9, CL 76:9, CL 82:11, CL 88:6, CL84:4 and CL 84:6 displayed a 2-fold increase in abundance within the NP treatment larvae (Fig. 6).

Within the energy storage lipids (i.e. glycerolipids, sterols, wax esters, hydroxy fatty acids, and free fatty acids), only 63 out of 278 lipid species displayed a difference in relative contribution by either not being present or being present in very low abundance in the Control compared to the NP treatments (Fig. S2). The heatmaps highlighted an increase in abundance, as well as a change in molecular speciation in some TAG, MADAG, MGDG, DGDG, OAHFA and FFA species; with a clear control versus NP treatment clustering in MADAG, OAFA and FFA (Fig. 7). Within the TAG species, TAG 38:0, 40:0, 42:0, 42:1, 42:2, 44:1, 44:2, and 46:2 displayed a 2-fold increase in abundance in both NP treatments compared with the Control. Of particular interest, 15 of 16 OAHFA species and two of five FFA species were not present in the Control, and the abundance of the remaining species belonging to both lipid classes increased with NP treatment concentration (Fig. 7).

**Figure 7.**
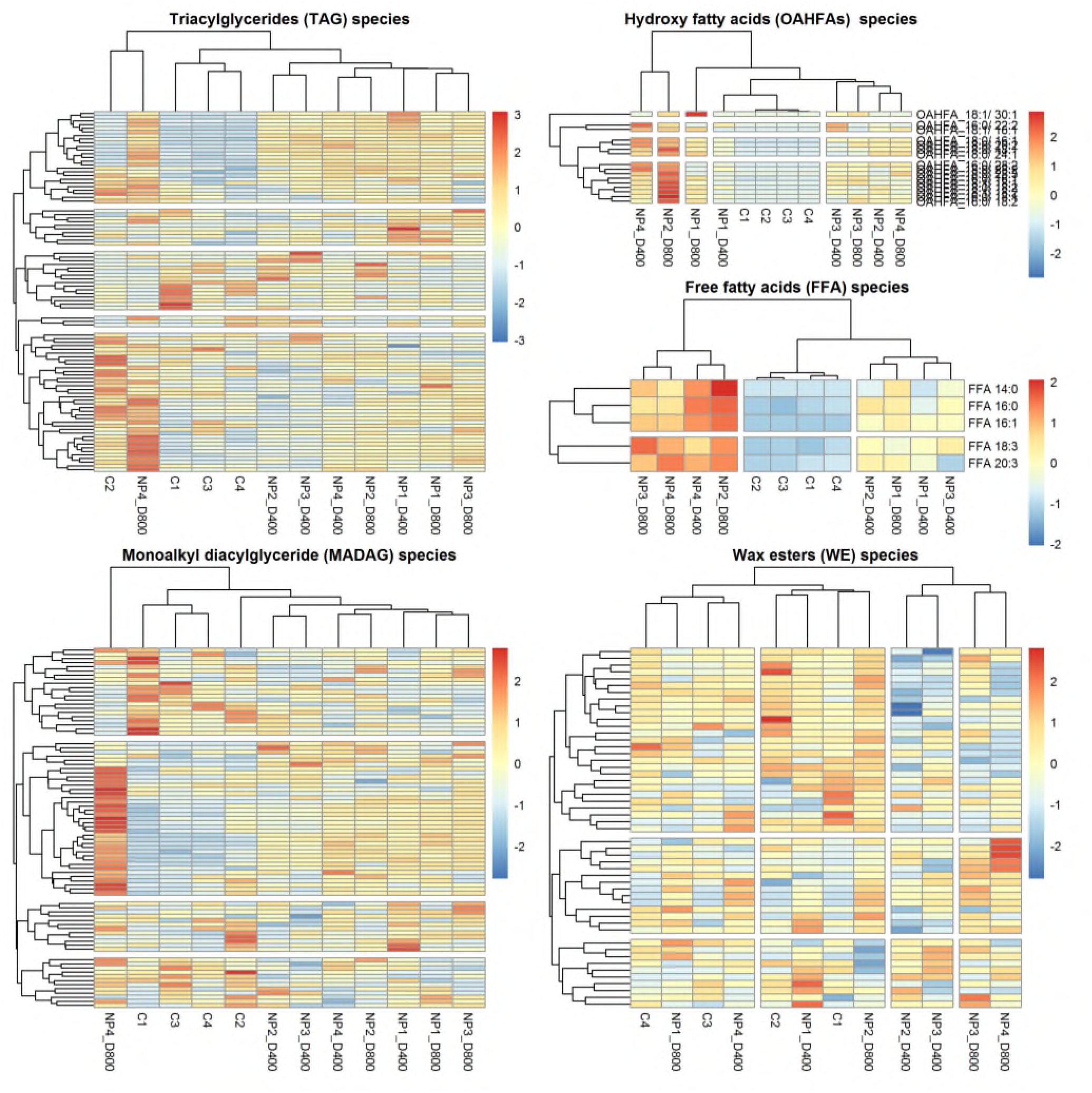
Heatmaps of glycerolipids molecular species within each lipid class detected in treatments (NP1 to NP4) and control (C1 to C4). Red and blue correspond to higher and lower relative amounts, respectively. Data were scaled by row.

## DISCUSSION

Coral larvae can consume up to 75% of their initial lipid reserves during their first week of life ^21,24,44^ and up to 2-fold post-settlement (Boulotte *et al.*, under review); therefore, depleted energy resources during the larval stage may play an essential role in post-settlement survival. Our study is the first attempt using an oil-in-water nanoemulsion to deliver marine triacylglycerides (TAG) to coral larvae with the aim to provide a nutritional boost to young corals. The provision of exogenous lipids during the larval stage resulted in an overall 30% increase in energetic lipids; which translated into a 46% increase in survival at 16 weeks post-settlement.

Coral larvae can obtain a considerable amount of energy from wax esters and TAG, with the former being slowly consumed during embryogenesis, while the latter can be rapidly hydrolysed for short-term energy needs ^21,26^. Samples for lipidomics analysis were taken eight hours post-nanoparticle exposure, which suggests that coral larvae can quickly hydrolyse TAG for short-term energy needs, as observed among other marine organisms (reviewed in ^21^). The results suggest that following larval feeding, depleted energy resources during the larval stage were no longer a limiting factor in post-settlement survival, highlighting that, nutritional experiences during the larval stage may influence the fitness of coral recruits.

The exposure to the nanoparticles triggered substantial and complex changes within the coral larvae lipidome composition via remodelling of both phospholipid and glycerolipid molecular species. Most changes occurred within the phospholipids, with the cardiolipin group (CL) being the most affected by the nanoparticle. CL is a signature lipid of the inner mitochondrial membrane and has been shown to play a significant role in mitochondrial bioenergetic processes and associated essential metabolic pathways ^45^. Therefore, the remodelling of CL observed post nanoparticle exposure, suggests that some bioenergetic processes may have occurred. Mitochondrial bioenergetics involves energy production via fatty acid oxidation, which starts with the hydrolysis of TAG into fatty acids (FAs) and glycerol, after which, FAs are transported into mitochondria for degradation and energy production, while the glycerol molecule is esterified in TAG or phospholipids ^46^. Therefore, we can speculate that 1) the exogenous TAG were hydrolysed into energy production in the larvae, and that, 2) the glycerol product of this hydrolysis was used to increase larval energetic reserves via *de novo* TAG synthesis; which corroborates with the increase levels in TAG along with changes in its molecular lipid species observed in the treatment larvae. Another result supporting TAG hydrolysis is the increase in free fatty acids in the treatment larvae, but also the presence of lysophosphatidylcholines (LPC), which are messenger lipids resulting from TAG hydrolysis ^47^. The increase in other phospholipid groups observed post nanoparticle exposure, may result from the esterified glycerol product of the TAG hydrolysis as glycerol is a known precursor for *de novo* phospholipids synthesis ^48^. Overall, our results highlight that *A.tenuis* larvae were able to ingest the nanoparticles and utilise the content for energy production and lipid storage.

Although coral larvae have been considered as non-feeding lecithotrophic larvae, there is a common misnomer about the dichotomy of planktotrophic vs lecithotrophic larvae despite many intermediate nutritional forms being described (reviewed in ^27^). A handful of studies from the 1980-90s reported observations of larval feeding among different coral species, such as Tranter et al., ^49^ who observed the larvae of ahermatypic coral, *Caryophyllia smithi*, developing mucus strings to trap particulate food. Other feeding behaviours were observed in the larvae of solitary coral *Fungia scutaria* ^50,51^, in *Cyphastra ocellina* ^52^, and in the soft coral *Heteroxenia fuscescens*, which were capable of uptaking dissolved free amino acids from the water column^53^. More recently, larvae from the cold-water coral, *Lophelia pertusa*, have been observed to feed by developing a protractible mouth 30 days after fertilisation ^54^. These observations are congruent with our results which indicate that coral larvae can ingest nanoparticles and absorb the lipid contents for immediate energy and storage. Therefore, we suggest that a continuum of nutritional strategies should be more widely recognised in coral early-life stages.

Most importantly, our results highlight that *A. tenuis* larvae should be considered as facultative feeders as part of their life history strategy. The ability to feed in the plankton provides benefits, as demonstrated by the uptake of lipids from the nanoparticles during the larval stage resulting in an increase in post-settlement survival. This supports the recognition of facultative feeding among other marine invertebrate larvae such as in the gastropod *Alderia* sp. ^55^, the polychaete annelid *Streblospio benedicti* ^56^, in echinoderms of the genus *Macrophiothrix* ^57^, and in the sea urchin *Tripneustes gratilla* larvae ^58^. The results from this study provide a paradigm shift in our understanding of the great diversity of nutritional patterns in marine invertebrate larvae, with practical applications for coral reef restoration based on sexual reproduction.

Solutions to improve the effectiveness of sexual coral propagation have gained momentum in the past decade, and research targeting the enhancement of early-life stages survival is now a high-priority in coral restoration interventions ^16^. Ecologically meaningful reef restoration requires restoration initiatives to be more cost-effective and undertaken at a large scale ^8^. Our innovative approach aimed at increasing coral post-settlement survival rate is simple, fast, and affordable. Therefore, assuming other coral species are facultative feeders, this method could be easily implemented by restoration practitioners to maximise coral restoration outcomes. However, additional research is required to verify nutritional enhancements of coral larvae across a range of coral species, different nanoparticle compositions, and longer post-settlement monitoring times. Furthermore, such advances in coral restoration initiatives will only be effective if immediate actions are implemented to reduce CO_2_ emissions to mitigate environmental changes.

## ACKNOWLEDGEMENTS

We thank A. Negri’s team from the Australian Institute of Marine Science (AIMS) and I. Clark for assisting with coral spawning and larval rearing. We also thank the AIMS SeaSim staff for logistical support, maintenance of coral recruits and survival monitoring photographs. N.M.B. acknowledges funding provided by Southern Cross University through a Marine Ecology Research Centre grant, and postgraduate research grant. P.L.H. acknowledges funding support provided by grants from the Australian Centre for International Agricultural Research (FIS 2014 063), and the Great Barrier Reef Foundation.

## AUTHOR CONTRIBUTION

N.M.B., D.R. and K.B. conceived and designed the study. N.M.B. and D.R. conducted the experiment. C.H. coordinated the settlement and recruit survival monitoring. N.M.B. analysed and interpreted all data. N.M.B. wrote the first manuscript draft, and all authors contributed to revising the drafts and final version.

