## Supplementary material for "Nanodelivery of lipids to coral larvae maximises post-settlement survival: Implications for larval ecology and reef restoration": Figure S1

### SUPPLEMENTARY INFORMATION

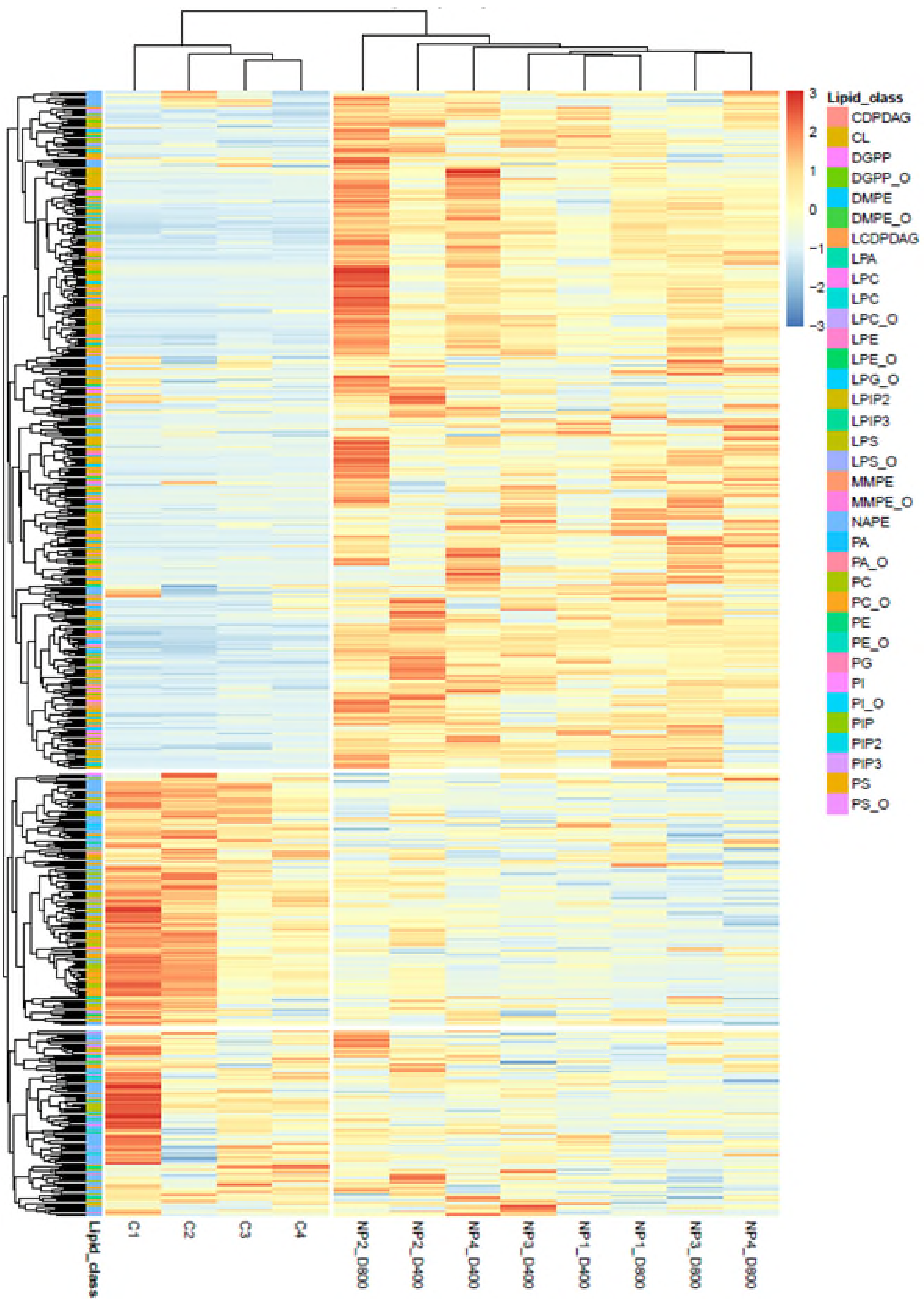


***Figure S1*** *Heatmaps of phospholipids molecular species within each lipid class detected in treatments (NP1 to NP4) and control (C1 to C4). Red and blue correspond to higher and lower relative amounts, respectively. Data were scaled by row.*

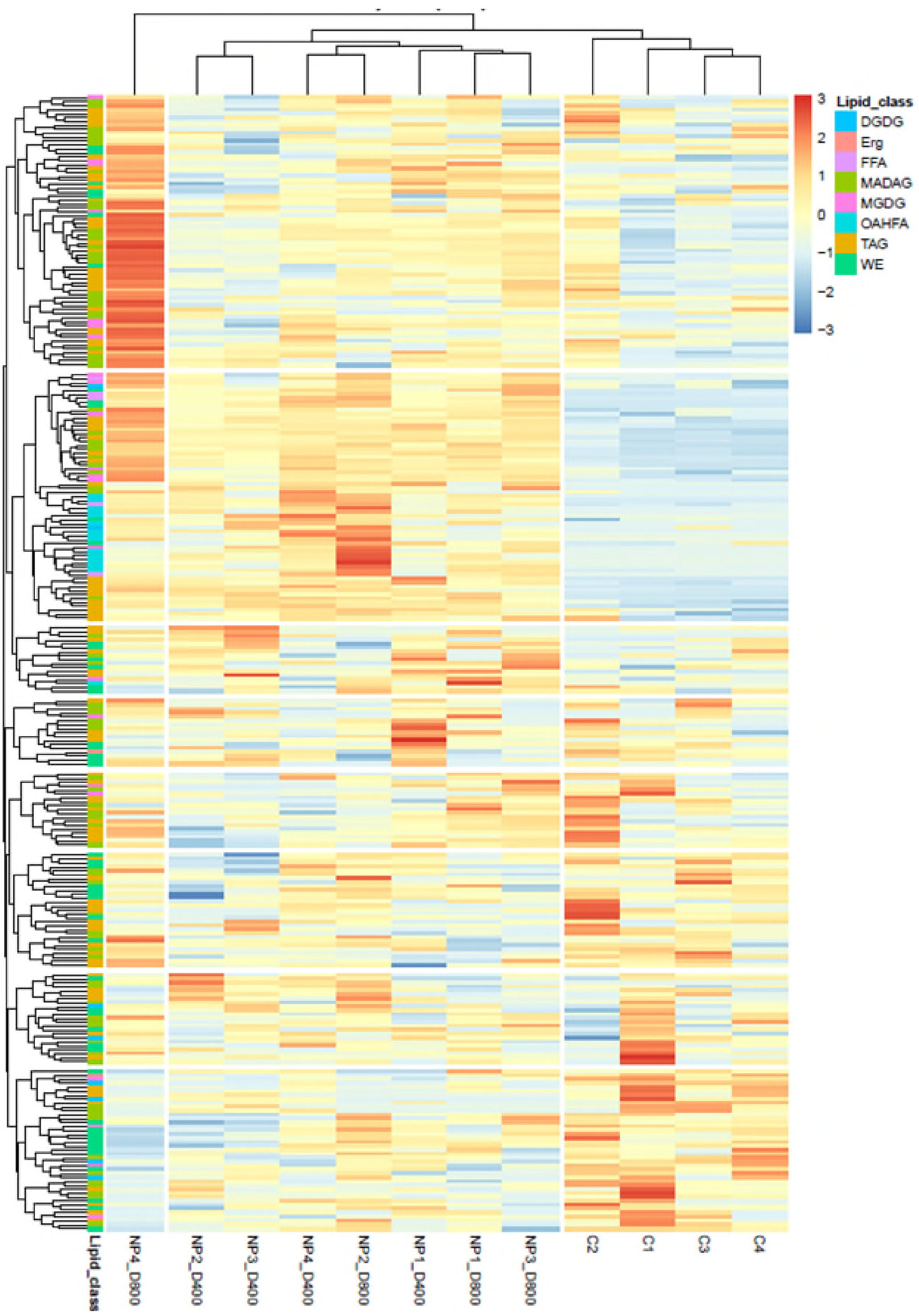


***Figure S2.*** *Heatmaps of glycerolipids molecular species within each lipid class detected in treatments (NP1 to NP4) and control (C1 to C4). Red and blue correspond to higher and lower relative amounts, respectively. Data were scaled by row.*

***Table S1. PERMANOVA results of the entire coral larvae lipidome composition***

| **Source** | **df** | **SS** | **MS** | **Pseudo-F** | **P(perm)** | **Unique perms** |
|---|---|---|---|---|---|---|
| **Tr** | 3 | 1113 | 371 | 17.819 | 0.0001 | 9930 |
| **Res** | 12 | 249.85 | 20.821 | | | |
| **Total** | 15 | 1362.9 | | | | |

*Details of the expected mean squares (EMS) for the model*

| **Source** | **EMS** |
|---|---|
| **Tr** | 1*V(Res) + 4*S(Tr) |
| **Res** | 1*V(Res) |

*Construction of Pseudo-F ratio(s) from mean squares*

| **Source** | **Numerator** | **Denominator** | **Num.df** | **Den.df** |
|---|---|---|---|---|
| **Tr** | 1*Tr | 1*Res | 3 | 12 |

*Estimates of components of variation*

| **Source** | **Estimate** | **Sq.root** |
|---|---|---|
| **S(Tr)** | 87.545 | 9.3566 |
| **V(Res)** | 20.821 | 4.563 |

***Table S2. Pair-wise tests results of the entire coral larvae lipidome composition***

| **Groups** | **t** | **P(perm)** | **Unique perms** |
|---|---|---|---|
| **Control, NP (400)** | 5.4039 | 0.0265 | 35 |
| **Control, NP (800)** | 5.6801 | 0.0305 | 35 |
| **Control, NP (0)** | 2.5249 | 0.0286 | 35 |
| **NP (400), NP (800)** | 0.94362 | 0.4802 | 35 |
| **NP (400), NP (0)** | 3.8458 | 0.0298 | 35 |
| **NP (800), NP (0)** | 4.1258 | 0.0316 | 35 |

*Denominators*

| **Groups** | **Denominator** | **Den.df** |
|---|---|---|
| Control, NP (400) | 1*Res | 6 |
| Control, NP (800) | 1*Res | 6 |
| Control, NP (0) | 1*Res | 6 |
| NP (400), NP (800) | 1*Res | 6 |
| NP (400), NP (0) | 1*Res | 6 |
| NP (800), NP (0) | 1*Res | 6 |

*Average Similarity between/within groups*

| | **Control** | **NP (400)** | **NP (800)** | **NP (0)** |
|---|---|---|---|---|
| **Control** | 92.264 | | | |
| **NP (400)** | 80.978 | 94.749 | | |
| **NP (800)** | 79.204 | 94.583 | 94.116 | |
| **NP (0)** | 89.61 | 88.018 | 86.508 | 93.946 |

***Table S3. PERMANOVA results within the energetic lipids class***

| Source | df | SS | MS | Pseudo-F | P(perm) | Unique perms |
|---|---|---|---|---|---|---|
| **Tr** | 2 | 284.75 | 142.38 | 17.815 | 0.0029 | 4743 |
| **Res** | 9 | 71.928 | 7.992 | | | |
| **Total** | 11 | 356.68 | | | | |

*Details of the expected mean squares (EMS) for the model*

| Source | EMS |
|---|---|
| **Tr** | 1*V(Res) + 4*S(Tr) |
| **Res** | 1*V(Res) |

*Construction of Pseudo-F ratio(s) from mean squares*

| Source | Numerator | Denominator | Num.df | Den.df |
|---|---|---|---|---|
| **Tr** | 1*Tr | 1*Res | 2 | 9 |

*Estimates of components of variation*

| Source | Estimate | Sq.root |
|---|---|---|
| **S(Tr)** | 33.596 | 5.7962 |
| **V(Res)** | 7.992 | 2.827 |

***Table S4. PERMANOVA results within the structural lipids class***

| Source | df | SS | MS | Pseudo-F | P(perm) | Unique perms |
|---|---|---|---|---|---|---|
| **Tr** | 2 | 888.77 | 444.39 | 31.885 | 0.0027 | 4763 |
| **Res** | 9 | 125.44 | 13.937 | | | |
| **Total** | 11 | 1014.2 | | | | |

*Details of the expected mean squares (EMS) for the model*

| Source | EMS |
|---|---|
| **Tr** | 1*V(Res) + 4*S(Tr) |
| **Res** | 1*V(Res) |

*Construction of Pseudo-F ratio(s) from mean squares*

| Source | Numerator | Denominator | Num.df | Den.df |
|---|---|---|---|---|
| **Tr** | 1*Tr | 1*Res | 2 | 9 |

*Estimates of components of variation*

| Source | Estimate | Sq.root |
|---|---|---|
| **S(Tr)** | 107.61 | 10.374 |
| **V(Res)** | 13.937 | 3.7333 |

***Table S5. PERMANOVA results within the fatty acids class***

| Source | df | SS | MS | Pseudo-F | P(perm) | Unique perms |
|---|---|---|---|---|---|---|
| **Tr** | 2 | 664.19 | 332.1 | 7.6766 | 0.0016 | 4762 |
| **Res** | 9 | 389.34 | 43.261 | | | |
| **Total** | 11 | 1053.5 | | | | |

*Details of the expected mean squares (EMS) for the model*

| Source | EMS |
|---|---|
| **Tr** | 1*V(Res) + 4*S(Tr) |
| **Res** | 1*V(Res) |

*Construction of Pseudo-F ratio(s) from mean squares*

| Source | Numerator | Denominator | Num.df | Den.df |
|---|---|---|---|---|
| **Tr** | 1*Tr | 1*Res | 2 | 9 |

*Estimates of components of variation*

| Source | Estimate | Sq.root |
|---|---|---|
| **S(Tr)** | 72.209 | 8.4976 |
| **V(Res)** | 43.261 | 6.5773 |

***Table S6. PERMANOVA results within the cell signalling lipids class***

| Source | df | SS | MS | Pseudo-F | P(perm) | Unique perms |
|---|---|---|---|---|---|---|
| **Tr** | 2 | 25.79 | 12.895 | 1.515 | 0.1254 | 4730 |
| **Res** | 9 | 76.603 | 8.5114 | | | |
| **Total** | 11 | 102.39 | | | | |

*Details of the expected mean squares (EMS) for the model*

| Source | EMS |
|---|---|
| **Tr** | 1*V(Res) + 4*S(Tr) |
| **Res** | 1*V(Res) |

*Construction of Pseudo-F ratio(s) from mean squares*

| Source | Numerator | Denominator | Num.df | Den.df |
|---|---|---|---|---|
| **Tr** | 1*Tr | 1*Res | 2 | 9 |

*Estimates of components of variation*

| Source | Estimate | Sq.root |
|---|---|---|
| **S(Tr)** | 1.0959 | 1.0469 |
| **V(Res)** | 8.5114 | 2.9174 |

*Table S7. SIMPER analysis results within the energetic lipids class*

***Groups Control & NP (400)***

Average dissimilarity = 10.27

| | Group Control | Group NP (400) | | | | |
|---|---|---|---|---|---|---|
| **Species** | **Av.Abund** | **Av.Abund** | **Av.Diss** | **Diss/SD** | **Contrib%** | **Cum.%** |
| **TAG** | 4.73 | 5.87 | 2.55 | 3.73 | 24.80 | 24.80 |
| **WE_16:0/** | 6.80 | 5.77 | 2.31 | 4.87 | 22.46 | 47.25 |
| **FFA** | 0.32 | 0.81 | 1.10 | 2.45 | 10.67 | 57.92 |
| **OAHFA_18:0/** | 0.00 | 0.39 | 0.87 | 3.68 | 8.47 | 66.39 |
| **OAHFA_16:0/** | 0.05 | 0.37 | 0.70 | 2.18 | 6.85 | 73.24 |

***Groups Control & NP (800)***

Average dissimilarity = 11.16

| | Group Control | Group NP (800) | | | | |
|---|---|---|---|---|---|---|
| **Species** | **Av.Abund** | **Av.Abund** | **Av.Diss** | **Diss/SD** | **Contrib%** | **Cum.%** |
| **WE_16:0/** | 6.80 | 5.60 | 2.68 | 3.03 | 23.98 | 23.98 |
| **TAG** | 4.73 | 5.77 | 2.32 | 2.97 | 20.83 | 44.81 |
| **FFA** | 0.32 | 0.91 | 1.31 | 3.52 | 11.74 | 56.55 |
| **OAHFA_18:0/** | 0.00 | 0.41 | 0.91 | 3.53 | 8.14 | 64.69 |
| **OAHFA_16:0/** | 0.05 | 0.42 | 0.82 | 2.66 | 7.34 | 72.03 |

*Table S8. SIMPER analysis results within the structural lipids class*

***Groups Control & NP (400)***

Average dissimilarity = 17.81

| | Group Control | Group NP (400) | | | | |
|---|---|---|---|---|---|---|
| **Species** | **Av.Abund** | **Av.Abund** | **Av.Diss** | **Diss/SD** | **Contrib%** | **Cum.%** |
| **CL** | 1.99 | 4.30 | 4.74 | 3.68 | 26.61 | 26.61 |
| **PC** | 7.62 | 6.55 | 2.20 | 3.38 | 12.34 | 38.94 |
| **PA** | 0.67 | 1.65 | 2.01 | 3.48 | 11.28 | 50.22 |
| **DGPP** | 0.14 | 0.80 | 1.34 | 2.36 | 7.51 | 57.74 |
| **PI** | 0.26 | 0.91 | 1.33 | 3.84 | 7.48 | 65.22 |
| **PS** | 2.64 | 2.06 | 1.19 | 3.11 | 6.67 | 71.89 |

***Groups Control & NP (800)***

Average dissimilarity = 19.17

| | Group Control | Group NP (800) | | | | |
|---|---|---|---|---|---|---|
| **Species** | **Av.Abund** | **Av.Abund** | **Av.Diss** | **Diss/SD** | **Contrib%** | **Cum.%** |
| **CL** | 1.99 | 4.28 | 4.67 | 3.50 | 24.35 | 24.35 |
| **PC** | 7.62 | 6.40 | 2.49 | 4.14 | 12.97 | 37.32 |
| **PA** | 0.67 | 1.80 | 2.30 | 3.56 | 11.99 | 49.31 |
| **PI** | 0.26 | 1.06 | 1.62 | 4.65 | 8.45 | 57.75 |
| **DGPP** | 0.14 | 0.92 | 1.57 | 3.89 | 8.21 | 65.96 |
| **MMPE** | 0.28 | 0.87 | 1.20 | 5.59 | 6.28 | 72.25 |

*Table S9. SIMPER analysis results within the fatty acid class*

***Groups Control & NP (400)***

Average dissimilarity = 15.21

| | Group Control | Group NP (400) | | | | |
|---|---|---|---|---|---|---|
| **Species** | **Av.Abund** | **Av.Abund** | **Av.Diss** | **Diss/SD** | **Contrib%** | **Cum.%** |
| **C10:0** | 2.14 | 3.27 | 1.76 | 1.44 | 11.60 | 11.60 |
| **C16:0** | 1.78 | 2.90 | 1.72 | 1.84 | 11.33 | 22.94 |
| **C12:0** | 0.00 | 1.12 | 1.72 | 4.56 | 11.31 | 34.24 |
| **C20:4** | 5.47 | 4.67 | 1.24 | 2.31 | 8.13 | 42.37 |
| **C18:0** | 4.82 | 4.08 | 1.17 | 1.62 | 7.70 | 50.07 |
| **C22:4** | 2.21 | 1.52 | 1.14 | 1.71 | 7.52 | 57.59 |
| **C20:1** | 0.37 | 1.10 | 1.13 | 1.74 | 7.40 | 64.99 |
| **C18:1** | 3.69 | 3.98 | 0.81 | 1.08 | 5.35 | 70.34 |

***Groups Control & NP (800)***

Average dissimilarity = 18.88

| | Group Control | Group NP (800) | | | | |
|---|---|---|---|---|---|---|
| **Species** | **Av.Abund** | **Av.Abund** | **Av.Diss** | **Diss/SD** | **Contrib%** | **Cum.%** |
| **C10:0** | 2.14 | 3.80 | 2.53 | 2.39 | 13.43 | 13.43 |
| **C16:0** | 1.78 | 3.25 | 2.24 | 2.80 | 11.88 | 25.31 |
| **C12:0** | 0.00 | 1.37 | 2.09 | 8.77 | 11.07 | 36.38 |
| **C20:4** | 5.47 | 4.17 | 1.98 | 2.41 | 10.50 | 46.88 |
| **C18:0** | 4.82 | 3.84 | 1.49 | 1.95 | 7.90 | 54.79 |
| **C14:0** | 0.65 | 1.44 | 1.20 | 5.31 | 6.33 | 61.12 |
| **C22:4** | 2.21 | 1.55 | 1.12 | 1.72 | 5.94 | 67.06 |
| **C20:5** | 3.23 | 2.67 | 1.09 | 1.73 | 5.79 | 72.85 |

*Table S10. List of abbreviated lipid classes*

| Lipid Class | Lipid code |
|---|---|
| Phosphatidic Acid | **PA** |
| Phosphatidylcholine | **PC** |
| Phosphatidylethanolamine | **PE** |
| Phosphatidylinositol | **PI** |
| Phosphatidylserine | **PS** |
| Cardiolipin | **CL** |
| Diacylglycerol Pyrophosphate | **DGPP** |
| Monomethyl-phosphatidylethanolamine | **MMPE** |
| Dimethyl-phosphatidylethanolamine | **DMPE** |
| Phosphatidylinositol | **PIP** |
| Phosphatidylinositol 4,5-bisphosphate | **PIP2** |
| Phosphatidylinositol (3,4,5)-trisphosphate | **PIP3** |
| Cytidine Diphosphate Diacylglycerol | **CDPDAG** |
| N-Acylphosphatidylethanolamine | **NAPE** |
| Sphingomyelin | **SM** |
| Ceramide | **Cer** |
| Sulphatides | **SGalCer** |
| Ceramide-phosphate | **CerP** |
| Ceramide phosphatidylethanolamine | **CerPE** |
| Inositolphosphorylceramide | **IPC** |
| Mannosyl-inositolPceramide | **MIPC** |
| Mannosyl-diinositolPceramide | **M(IP)2C** |
| Trihexosylceramide | **Hex3Cer** |
| Monosialoganglioside | **GM3** |
| Ceramide Ciliatine | **CerCil** |
| Triacylglyceride | **TAG** |
| Monoalkyl diacylglyceride | **MADAG** |
| Monogalactosyl diacylglyceride | **MGDG** |
| Diagalactosyl diacylglyceride | **DGDG** |
| Free Fatty Acid | **FFA** |
| Wax Ester 16:0 | **WE_16:0/** |
| Wax Ester 18:0 | **WE_18:0/** |
